# Virus infection of honey bee queens alters lipid profiles and indirectly suppresses a retinue pheromone component via reducing ovary mass

**DOI:** 10.1101/2025.02.24.639811

**Authors:** Alison McAfee, Abigail Chapman, Armando Alcazar Magaña, Katie Marshall, Shelley E. Hoover, David R. Tarpy, Leonard J. Foster

## Abstract

Virus infections reduce honey bee (*Apis mellifera*) ovary mass (in part due to resource-allocation trade-offs with immunity) and are linked to the presence of supersedure cells in colonies, indicating that workers are attempting to replace their queen. Pheromones, lipids, and lipid transport proteins could mediate relationships among virus infection, ovary mass, and supersedure. When we infected honey bee queens in the laboratory and profiled their queen retinue pheromone (QRP) components from head extracts, we found that virus infections specifically reduced the QRP component methyl oleate. Data from an observational field study were consistent with this pattern. Lipidomics analysis of the same extracts suggests that virus infection decreases triacylglycerol abundances (major sources of stored energy). Reducing ovary mass via laying restriction was sufficient to lower methyl oleate abundance ― suggesting that virus infection reduces methyl oleate indirectly via ovary effects ― but was insufficient to reduce abundance of most triacylglycerols or stimulate immune effector expression. Abundance of circulating apolipophorin-III, a lipid transporter and putative potentiator of immune effectors, was lower in queens with restricted laying, suggesting that its expression may be controlled by nutrient availability while its immune-stimulating capacity is governed by other mechanisms. Prior research has shown that queen pheromone blends lacking methyl oleate are less attractive to workers; therefore, diminishing methyl oleate could result in a less desirable pheromone bouquet and possibly stimulate supersedure. The mechanism of methyl oleate reduction is yet to be determined, but is clearly tied to ovary size and possibly resource availability or mobilization.

## Introduction

As the sole reproductive female in a colony of tens of thousands, the honey bee (*Apis mellifera*) queen has an enormous reproductive output ― a feat that is enabled by the nutritious diet supplied to her by workers (facultatively sterile females) in addition to physiological and behavioral specialization (reviewed by Fèvre *et al.*^1^). During the seasonal period of peak colony growth (normally late spring), the queen lays approximately 850 to 3200 eggs per day, the weight of which are, on average, greater than her own body^2–4^. Such reproductive division of labor, one of the hallmarks of eusociality^5^, means that the queen does very little in the colony other than walk, lay eggs, and eat royal jelly provided by the workers. This jelly secretion ― originating from the workers’ hypopharyngeal and mandibular glands^6^― is rich in proteins, lipids, and carbohydrates^7,8^, providing the queen with all the resources required to enable nearly continuous egg laying.

Each mature oocyte is approximately 1.4 – 1.6 mm long^9^ and is provisioned with sufficient nutrients during oogenesis, which is carefully regulated by molecular “checkpoints”^10^, to enable the embryo to develop over the subsequent 72 hours post-laying. Assuming similarity to other insects, approximately 30-40% of the dry mass of oocytes is comprised of lipids^11^, nearly all of which are originally supplied to the queen in the form of fatty acids within royal jelly (in addition to what may be biosynthesized from carbohydrates in the queen’s fat body)^8,12^. Metabolic pathways and, in particular, lipid trafficking in queens, are therefore expected to be highly specialized processes that can support rapid turnover from nutrient inputs to egg outputs.

Provisions can enter oocytes developing in any of the queen’s ∼360 ovarioles^13^ by at least three distinct mechanisms: nurse cell dumping of cytoplasmic contents, receptor-mediated endocytosis, and enzyme-mediated delivery^11,14^, with lipids primarily entering via the latter two routes as lipoprotein cargo^11^. Vitellogenin, apolipophorin-III (ApoLP-III), and apolipophorins-I/II (ApoLP-I/II) are the major lipoproteins in insects^15,16^, and while they can all function as lipid carriers, vitellogenin and ApoLP-III are also pathogen-associated molecular pattern (PAMP) binding proteins with roles in insect immunity^17–19^. Vitellogenin’s main immunological role is to facilitate transgenerational immune priming by binding and transporting PAMPs into oocytes^20^, whereas ApoLP-III appears to act as a humoral immunity potentiator (reviewed by Weers and Ryan^15^). Vitellogenin is an inefficient lipid carrier (on a per protein basis), containing only 8- 10% lipids in its loaded form^11^, and is responsible for transferring only about 5% of the oocyte’s total lipid content^21^ from the hemolymph to the oocyte via receptor-mediated endocytosis^22^. Lipid-loaded ApoLP-I/II is a high-density lipoprotein (HDL; 30-50% lipid) and can also be taken up by oocytes via receptor-mediated endocytosis^11^; however, when ApoLP-I/II associates with loaded ApoLP-III particles, it forms a low-density lipoprotein (LDL) complex. These LDLs are responsible for delivering most (∼90%) of oocytes’ lipids, where entry occurs via enzyme-mediated delivery (in which the lipids, but not the protein, are taken up by the oocyte)^21^.

Though they are long-lived reproduction specialists, honey bee queens are still susceptible to many of the same pathogens that are abundant in adult workers, including viruses^23,24^. In our previous research, we found that virus infections can cause a reduction in the queens’ ovary mass (*i.e.,* a small-ovary phenotype) and compromise their ability to lay eggs^24,25^. Since immune stimulation alone was sufficient to reduce ovary mass, this implies that a resource allocation trade-off between reproduction and immunity underlies, at least in part, the small-ovary phenotype^25^; however, the mechanism of such a trade-off is still unknown, as is whether the trade-off occurs in a bi-directional manner (*i.e.*, is a reduction in ovary mass sufficient to increase investment in immunity?). Although many studies in insects have shown that increased fecundity is linked to poorer immunity (reviewed by Schwenke *et al.*^26^), the nature of this relationship is often complicated by life stage transitions that confound with fecundity^27–34^, and reproductive investment has not been experimentally reduced to test for reversibility.

Interestingly, ApoLP-III could mediate a reproduction-immunity trade-off, since some research in other insect species suggests its immune-stimulating and lipid transport functions are mutually exclusive^35^, and it appears to only function as an immune stimulator in its lipid-bound conformation^36–38^. We speculate that, upon associating with PAMPs, ApoLP-III may be unable to participate in HDL complexation and thus stymie the main route of lipid entry to oocytes. Likewise, when ovary investment is restricted and fewer lipids are being shuttled to oocytes, this may make more non-complexed ApoLP-III particles available to bind PAMPs. These ideas remain to be tested.

Over time, a honey bee queen’s fecundity does eventually wane, at which time the workers will initiate supersedure cell rearing to replace the existing queen^39^. The mechanism by which workers recognize and commit to this task is not known, but since previous research has identified correlations between queen fertility characteristics and pheromone abundance^40–44^, changes in queen pheromones could conceivably be involved. The honey bee queen retinue pheromone (QRP) is a synergistic blend of at least nine distinct compounds (fatty acids, esters, and alcohols) that elicit numerous and profound physiological and behavioral effects on drones (males) and workers^45,46^. Two competing theoretical frameworks explaining how such effects are realized are the “queen control hypothesis” (whereby the queen manipulates the workers via her pheromones) and the “honest signal hypothesis” (whereby pheromones convey information about the queen’s quality, allowing workers to evaluate the benefit of continued investment in her care)^47^. Both scenarios are compatible with a change in pheromonal signalling mediating the initiation of supersedure to some degree ― a decline in one or more pheromones or their components could either release the workers of queen control, or communicate to workers that the queen’s quality is no longer sufficient to support the colony’s population.

We previously observed that colonies headed by virus-infected queens were more likely to rear supersedure cells^25^, and here, we used a combination of newly-acquired and previously-published lipidomics^43^ and proteomics^48^ data to improve our understanding of the systems underlying virus infection and ovary investment. We used an established method^43^ to conduct untargeted lipidomics analysis on head extracts from these queens to test the hypothesis that virus infection is associated with a decline in one or more QRP components, thus providing a candidate mechanism for the observed supersedure cell rearing. Given the lipid demands of egg laying^11^ and that virus infection interferes with ovary investment^25^, we expected that virus infection would also be associated with broad lipidomic changes, and the untargeted nature of these data enabled us to explore such patterns. Because virus infection reduces queen ovary mass, a change in pheromone abundance could be a direct result of infection or an indirect ovary effect; therefore, to disentangle these relationships, we capitalized on previously published lipidomics data^43^ to evaluate QRP profiles in queens whose ovary masses were manipulated via laying restriction. Analysis of publicly available immune effector and major lipoprotein (vitellogenin, ApoLP-III, and ApoLP-I/II) data from these queens^48^ enabled us to test if the reproduction-immunity trade-off we previously observed functions in reverse ― that is, whether restricting ovary mass is sufficient to boost immune effector abundance ― and if abundance of circulating ApoLP-III is consistent with a potential role as a mediator.

## Results

### Effects of virus infection on QRP profiles and the lipidome

We measured seven components of the queen retinue pheromone (QRP) blend in queen head extracts and found that, among queens with experimentally manipulated virus levels in our previously published cage trial^25^, methyl oleate was the only QRP compound significantly linked to virus infection (F_1,24_ = 17.7, estimate = −0.33, p = 0.00031, α = 0.0071, Bonferroni correction; **Figure 1A**). Methyl oleate was also positively associated with ovary mass (though the relationship was marginally not significant; F_1,25_ = 6.3, estimate = 0.08, p = 0.019, α = 0.0071, Bonferroni correction; **Figure 1B**), as expected given that we previously established that queen viral load and ovary mass are inversely related^24,25^. Among queens heading colonies in the field, no QRP components were significantly linked to viral load, but one compound, again methyl oleate, was significantly positively linked to ovary mass (F_1,27_ = 10.2, estimate = 0.032, p = 0.0035, α = 0.0071, Bonferroni correction; **Figure 1C & D**).

**Figure 1.**
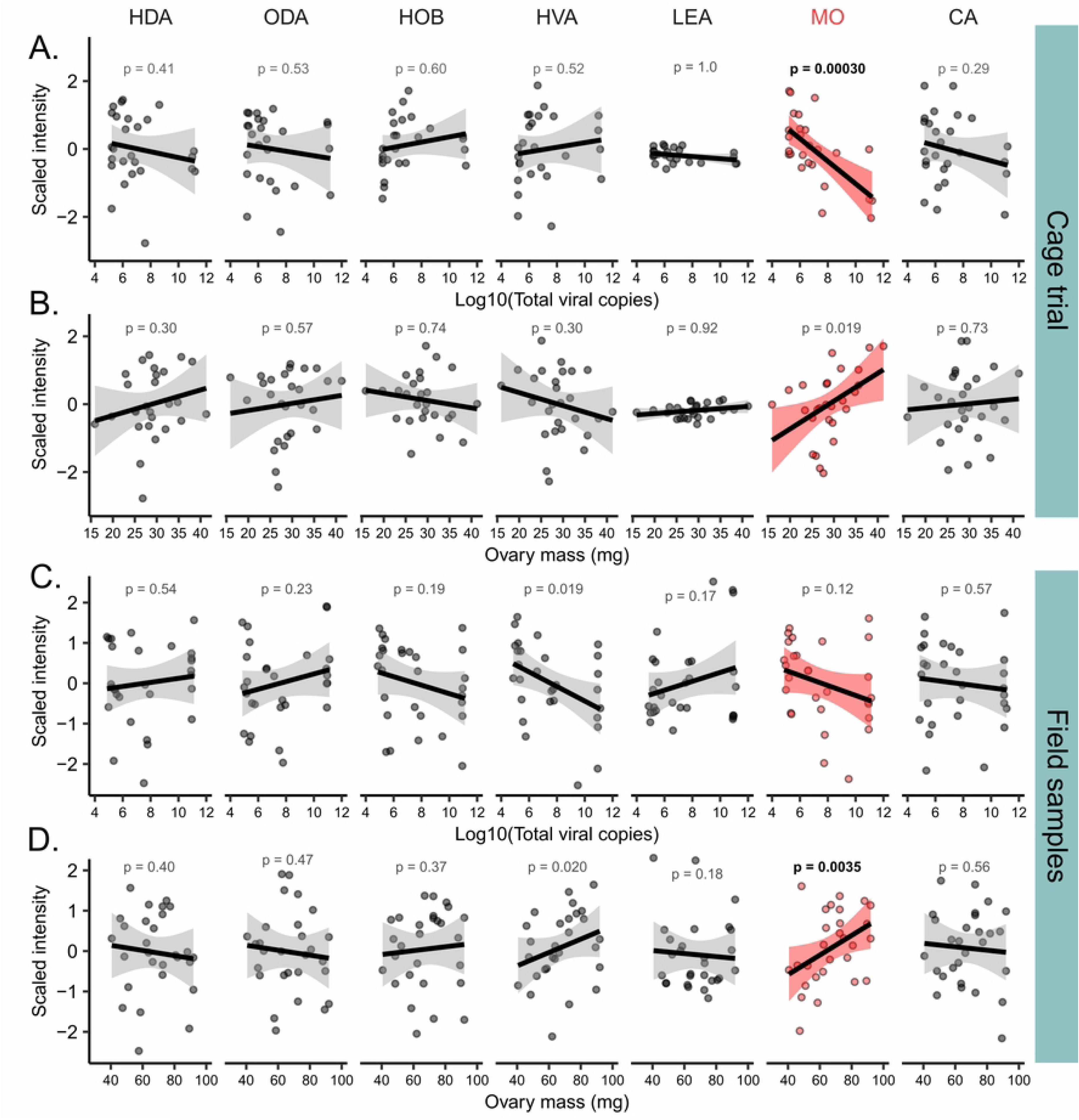
The QRP component methyl oleate is linked to queen virus infection and ovary mass. HDA = 9(R)-hydroxydec-2(E)-enoic acid; ODA = E-9-oxodec-2-enoic acid; HOB = methyl p-hydroxybenzoate; HVA = 4-hydroxy-3-methoxyphenylethanol; LEA = linolenic acid; MO = methyl oleate; CA = E-3-(4-hydroxy-3- methoxyphenyl)-prop-2-en-1-ol. Methyl oleate (red) was the only compound exhibiting significant correlations with tested variables (α = 0.0071; Bonferroni correction on 7 tests, bold text indicates p < α). Data were modelled using simple linear regressions. A) Compound intensities in relation to total viral load (cage trial). B) Compound intensities in relation to ovary mass (cage trial). C) Compound intensities in relation to total viral load (field samples). D) Compound intensities in relation to ovary mass (field samples).

To assess broader lipidomic changes related to queen virus infection, we conducted differential abundance testing of all annotated lipids evaluated in queen head extracts from the cage trial and field samples. Across both datasets, we identified 1,993 high-quality molecular features, of which 336 were annotated lipids, and 41 (12%) were triacylglycerols (**Supplementary Data 1**). Principal component (PC) analysis shows that samples weakly cluster according to total viral load in the cage trial, as indicated both visually (**Figure 2A**) and by regressing total viral load against PC1 (estimate = −0.070, t = −1.9, p = 0.071) and PC2 (estimate = −0.098, t = −1.7, p = 0.094). No interaction between PC1 and PC2 was found, and the whole model (total virus ∼ PC1+PC2) was marginally non-significant (F_2,22_ = 3.3, p = 0.055). The field samples, however, clustered more strongly according to total viral load (**Figure 2B)**, with both PC1 (estimate = −0.16, t = −4.2, p = 0.00029) and PC2 (estimate = 0.12, t = 2.8, p = 0.010) being significant predictors using the same model structure (whole model: F_2,26_ = 12.6, p = 0.00015; again no interaction was detected). Reflecting the PC analysis, only 4 out of 336 annotated lipids were significantly linked to total viral load among the cage trial queens, including methyl oleate as previously mentioned (FDR = 5%, Benjamini-Hochberg correction; **Figure 3A**). The additional lipids were all triacylglycerols (glyceryl trioleate, TG 18:1_18:1_20:1, and TG 18:0_19:1_21:1), and all were negatively linked to viral load. In the queens sampled from the field, a larger number (62) of annotated lipids were significantly linked to viral load (FDR = 5%, Benjamini-Hochberg correction; **Figure 3B**). Among these, only three increased with viral load, including a prostaglandin (PGG2), phosphatidylethanol (PEtOH 16:1_16:1), and sphingomyelin (SM 34:0;2O). All other compounds were downregulated with virus abundance, including 25 (61%) of the annotated triacylglycerols, and structural enrichment analysis shows that triacylglycerols were the most significantly enriched chemical class (**Figure 3C**). Pathway enrichment analysis was unfortunately not possible because exceedingly few (<10%) annotated lipids were associated with KEGG (Kyoto Encyclopedia of Genes and Genomes) identifiers.

**Figure 2.**
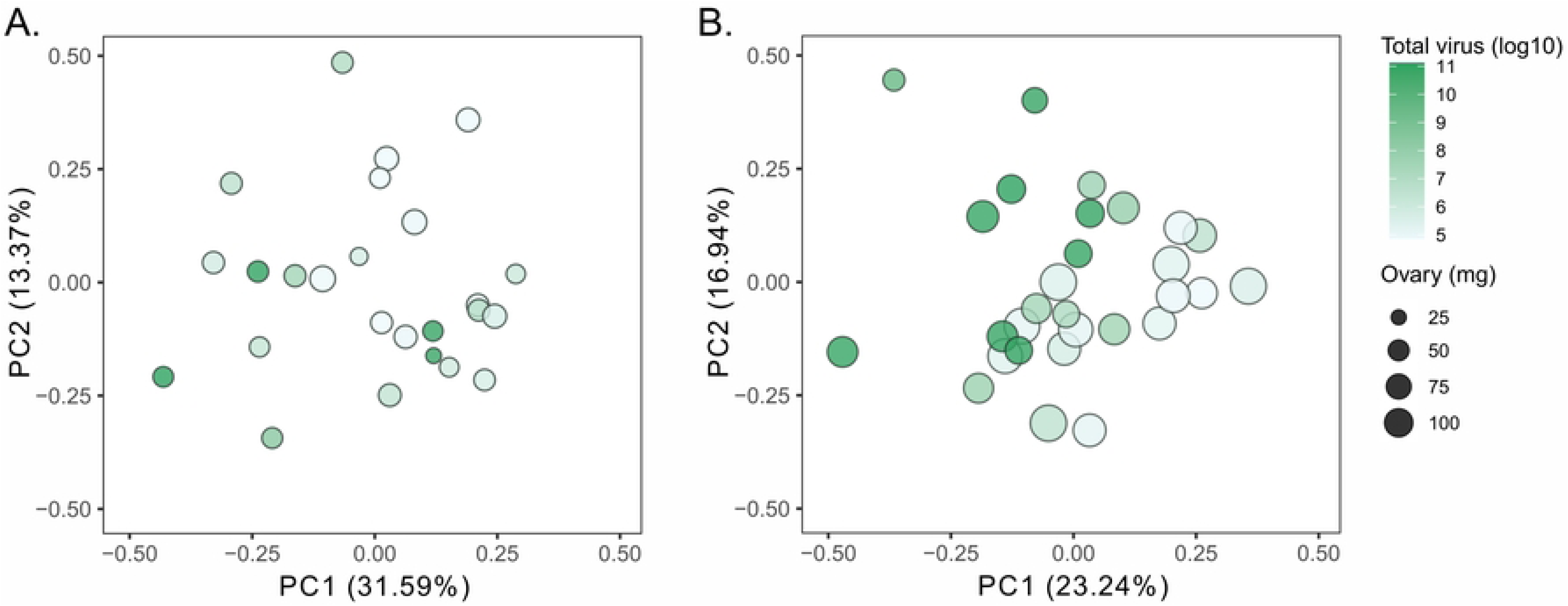
Principle component analysis suggests that lipid profiles are linked to viral loads in the field. Lipidomics analysis of queen head extracts identified 337 annotated compounds overall. A) Cage trial data. B) Field sample data. As previously reported ^25^, ovary masses among cage trial queens were significantly smaller than field trial queens due to reduced egg laying ability in the miniature laboratory laying cages.

**Figure 3.**
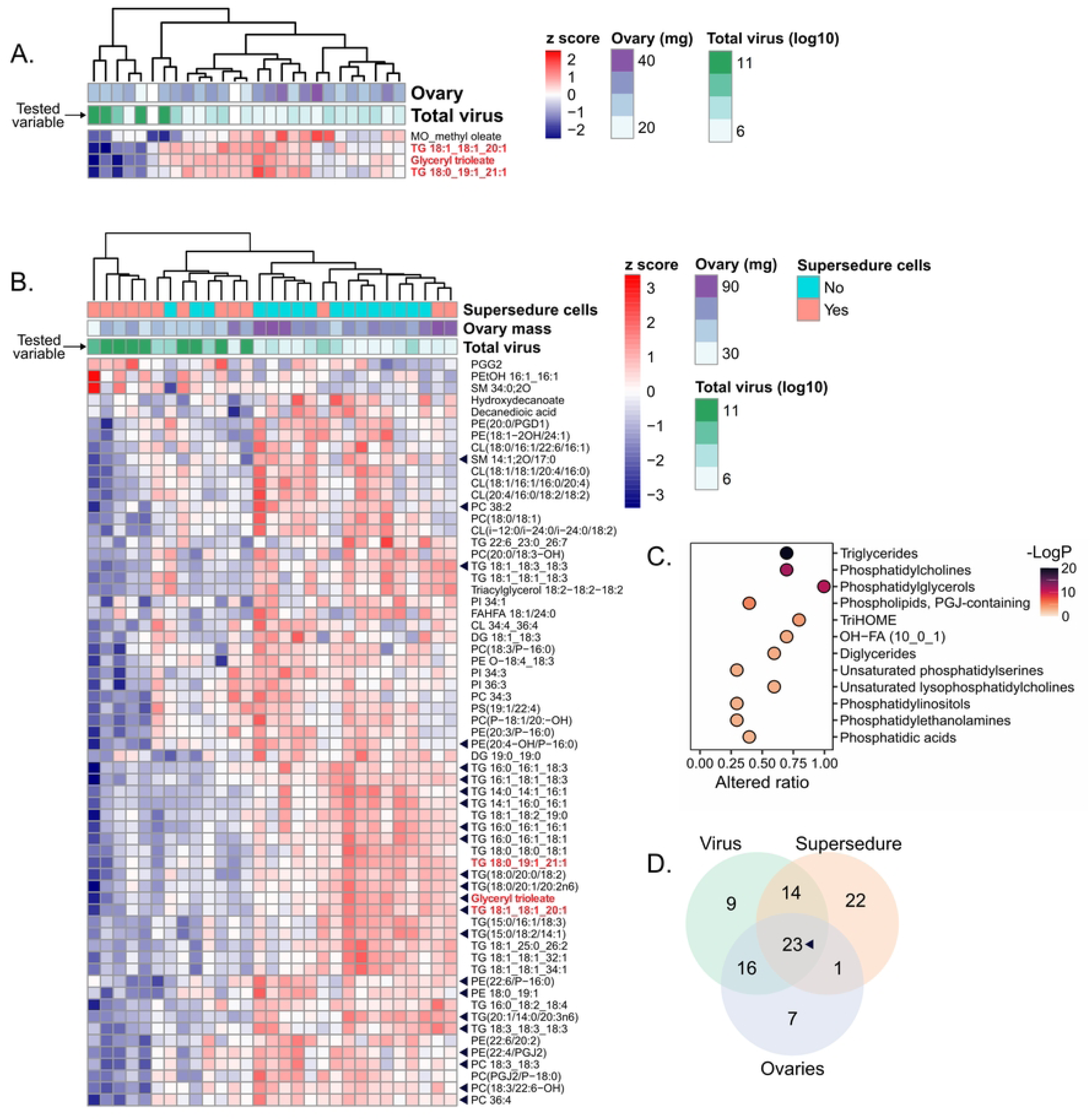
Annotated lipids significantly linked to virus infection. False discovery rates were controlled to 5% FDR (Benjamini-Hochberg correction). Differentially abundant compounds that are significant in both datasets (cage trial and field samples) are shown in red. A) 4 out of 336 annotated compounds were significantly linked to virus infection in the cage trial data. B) 62 compounds were significantly linked to virus infection in the field samples, 23 of which (black triangles) were also significantly linked to ovary mass and the presence of supersedure cells. C) Significantly enriched structural classes (5% FDR, Benjamini-Hochberg correction) in the field sample data. D) Overlap of annotated lipids significantly linked to total viral load, presence of supersedure cells, and ovary mass.

We previously identified a positive relationship between the viral load of queens and the presence of supersedure cells within colonies^25^, among other variables (see **Supplementary Figure S1**). Given the interrelatedness of queen total viral load, ovary mass, and the presence of supersedure cells, we analyzed the field-sampled queens’ lipidomic data with respect to each variable independently and identified overlapping differential abundance among comparisons. 23 lipids were significantly linked to all three variables (**Figure 3D**), of which 14 were triacylglycerols (34% of those identified), and two were among the four that were negatively linked to viral load in the cage trial queens (glyceryl trioleate and TG 18:1_18:1_20:1).

### QRP profiles and lipidomics of ovary-restricted queens

Larger colonies have queens with larger ovaries (**Figure S2**), but many confounding factors make the causal direction of this relationship uncertain. One way to manipulate ovary mass independent of other variables is to restrict ovary size via within-colony caging of otherwise equivalent queens^49^. We previously used this technique to produce queens with small (caged) and large (uncaged) ovaries^43^, and here we capitalized on those samples to determine what lipids, including methyl oleate, are specifically linked to ovary mass. Lipidomics analysis on head extracts from small- and large-ovary queens identified 1,739 molecular features, of which 250 were annotated, including only 13 (5.2%) triacylglycerols (**Supplementary Data 1**). 72 annotated lipids were differentially abundant in small-versus large-ovary queens (5% FDR, Benjamini-Hochberg method; **Figure S3**), including methyl oleate, which was higher in large-ovary queens, as predicted (**Figure 4A**). Only four triacylglycerols were differentially abundant, representing 31% of those identified, approximately half the fraction significantly linked to virus infection in the virus field trial queens. Two triacylglycerols were upregulated and two were downregulated, in contrast to the concerted downregulation of triacylglycerols with virus infection. Only 52 annotated lipids were detected in both the ovary restriction dataset and the field sample dataset described above, of which 19 (mostly phosphatidylethanolamines and phosphatidylinositols, and no triacylglycerols) were differentially abundant (**Figure 4B**). Ovary mass restriction via caging is thus sufficient to reduce methyl oleate abundance in addition to changing other lipids (**Figure S3**), but poor dataset overlap hampered our ability to compare ovary restriction versus virus effects on the lipidome.

**Figure 4.**
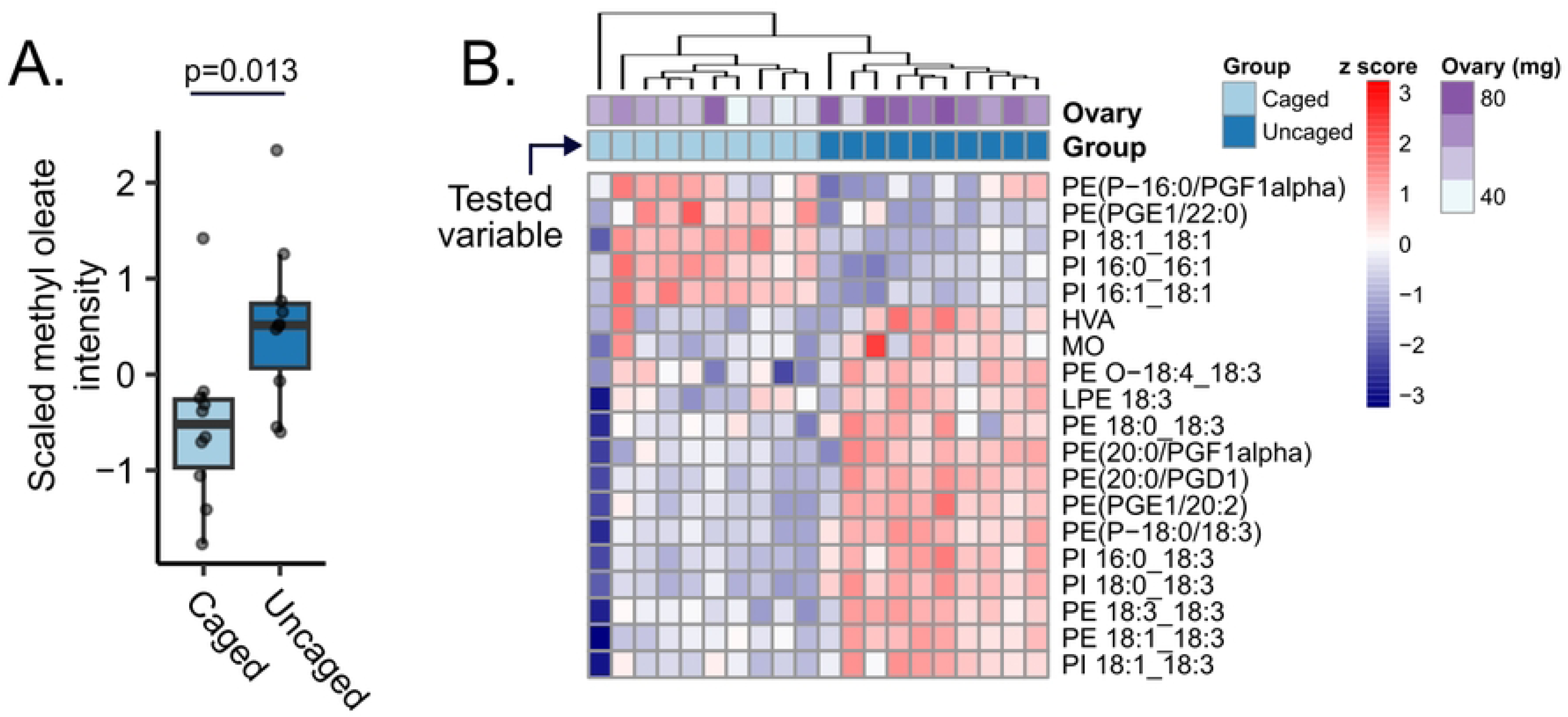
Ovary restriction is sufficient to reduce methyl oleate abundance. We used our previously published lipidomics dataset ^43^ of in-colony caged (small ovary) and uncaged (large ovary) queens to examine how lipids are affected by ovary restriction. A) Scaled methyl oleate abundance in caged and uncaged queens, analyzed using a simple linear model. B) Of the prior 336 lipids annotated in the field samples, 52 were also evaluated in the ovary restriction dataset and 19 of these (shown) were differentially abundant in small- vs. large-ovary queens among a larger pool of 72 differentially abundant annotated lipids (see **Figure S2** for the complete list). Benjamini-Hochberg correction, 5% FDR.

### Immune protein and lipid cargo protein expression

We previously found that the reduction in ovary mass caused by virus infection is in part a consequence of the cost of immune induction, rather than a direct pathological effect^25^. To test if this trade-off can operate in a bi-directional manner ― that is, if ovary restriction alone is sufficient to conversely elevate immune protein abundance ― we assessed immune proteins quantified in the hemolymph of the small- and large-ovary queens^48^. We quantified 3,164 unique protein groups in the hemolymph proteome, including all of the immune proteins that we previously found to be associated with virus infection^25^, but none significantly varied according to ovary size (5% FDR, Benjamini-Hochberg method; **Figure 5A, Figure S4**). A reduction in ovary mass is therefore not sufficient to enable induction of immune protein expression.

**Figure 5.**
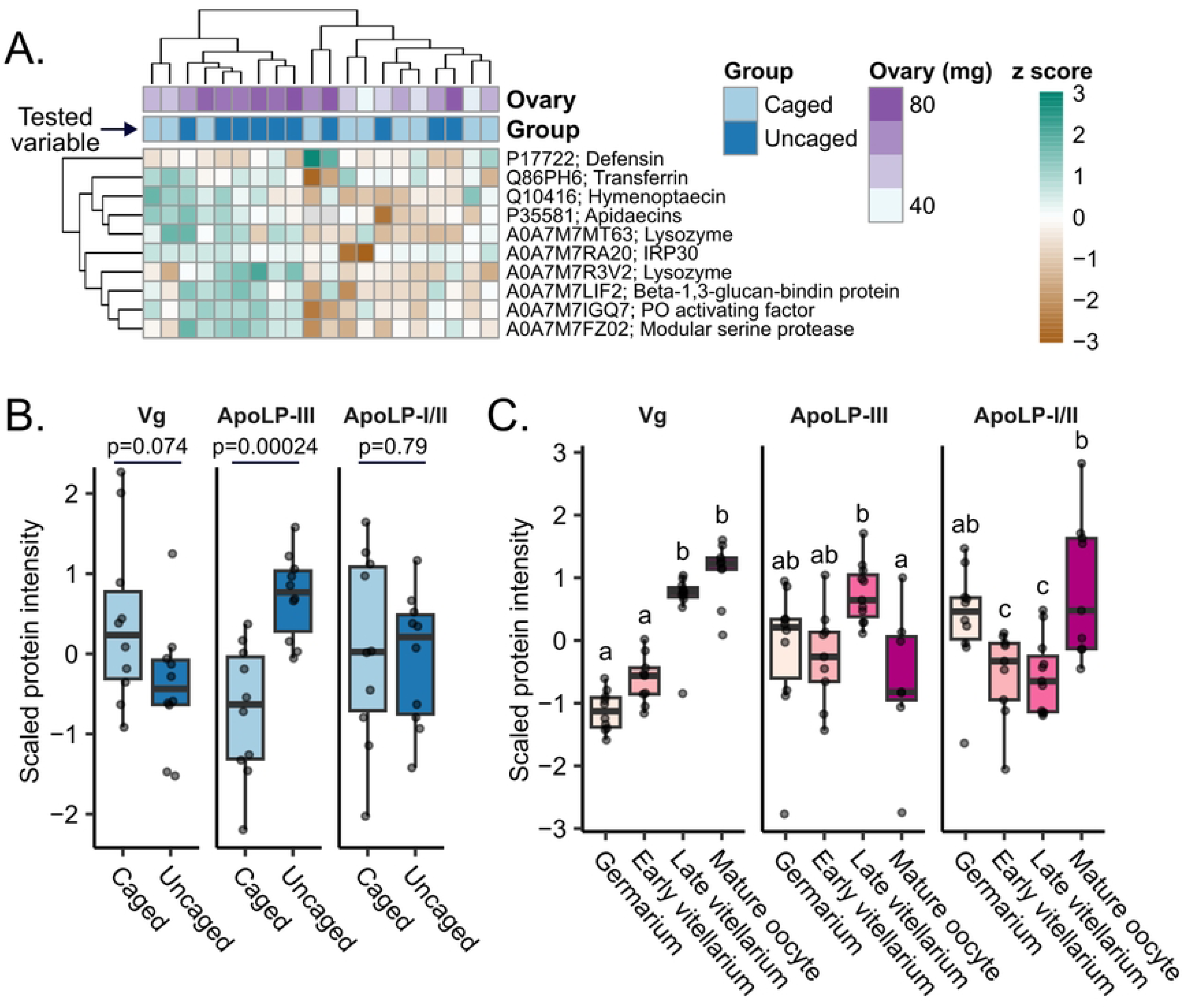
Ovary restriction affects ApoLP-III but is not sufficient to alter expression of circulating immune effectors. A) Immune effectors that were previously found to be stimulated by virus infection ^25^ are not significantly linked to small- (caged) vs. large-ovary (uncaged) queen groups (5% FDR, Benjamini-Hochberg correction). B) Analysis of specific circulating lipid transporters quantified in hemolymph of small- vs. large-ovary queens. P values indicate results of t-tests. Vg = vitellogenin; ApoLP-III = apolipophorin-III-like protein; ApoLP-I/II = apolipophorins-I/II. C) Lipid transporters quantified in progressive ovary sections (approximating the germarium, early vitellarium, and late vitellarium) and mature oocytes collected from the lateral oviduct. Different letters indicate statistical significance between groups, as determined by linear mixed modelling with section as a fixed effect (4 levels) and queen as a random effect (10 levels). Pairwise comparison p values were corrected by the Tukey method.

Apolipophorin (ApoLP)-III and ApoLP-I/II are major lipid cargo proteins, and ApoLP-III is especially interesting because it functions as a lipid transporter, pathogen recognition protein, and potentiator of immune effectors (see review^19^). Given their potential roles as a molecular switch system, we investigated circulating (hemolymph) ApoLP-III (Uniprot accession: B0LUE8) and ApoLP-I/II (A0A7M7SQ18) abundance, as well as the well-known lipoprotein vitellogenin (Q868N5) in small- and large-ovary queens. We found that ApoLP-III abundance was lower in queens with small ovaries (F_1,18_ = 20.9, estimate = −1.43, p = 0.00024; **Figure 5B**), despite immune effectors being unaltered, suggesting that the immune induction activity of ApoLP-III could be attenuated by reducing expression under conditions when resource availability is low. ApoLP-I/II did not differ (F_1,18_ = 0.076, estimate = −0.13, p = 0.79), and circulating vitellogenin slightly, but not significantly, declined in queens with larger ovaries (F_1,18_ = 3.6, estimate = −0.80, p = 0.074).

To investigate patterns of lipid transporter expression within ovaries, we analyzed three ovary sections approximately corresponding to the germarium (anterior section), early vitellarium (mid-ovary section), and late vitellarium (posterior section) in addition to mature eggs that we sampled from the oviducts. We found that while vitellogenin showed sequentially higher abundance in each successive section (the expected pattern based on progressive accumulation by receptor-mediated endocytosis), ApoLPs did not (**Figure 5C**). ApoLP-III was significantly less abundant in oocytes compared to late vitellarium sections (linear mixed model, estimate = −1.4, Kenward-Roger df estimate = 27.1, p = 0.012), which is consistent with enzyme-mediated delivery of cargo to oocytes (whereby the lipoprotein may still localize to the oocyte surface within ovarioles but is not endocytosed into the egg^11^). In contrast, ApoLP-I/II abundance increased in mature oocytes (estimate = 1.3, Kenward-Roger df estimate = 26.4, p = 0.0018; **Figure 5C**), suggesting that it can become endocytosed and concentrated in eggs. Major lipid trafficking systems in honey bee queens thus appear to be consistent with what is known in other insects.

## Discussion

Here we show that virus infection reduces the QRP component methyl oleate in queen heads (**Figure 1**), which offers a candidate mechanism for how workers may perceive that a queen has become infected and initiate supersedure. Since ovary restriction alone was sufficient to also reduce methyl oleate abundance (**Figure 4**), it may not be the infection, per se, that the workers are sensing; rather, it is more likely the change in ovary investment. The QRP component homovanillyl alcohol (HVA, or 4-hydroxy-3- methoxyphenylethanol) also declines with ovary mass^43^ and tended to decrease with virus infection in this study, but only in the field-sampled queens (and this trend was not significant after multiple hypothesis testing correction), whereas the patterns of methyl oleate abundance were more consistent. Further experimentation is needed to determine if a decline in methyl oleate is sufficient to trigger supersedure cell rearing, and whether it is based on absolute or relative pheromone quantities, but it is an intriguing and plausible hypothesis.

Virus infection was linked to broad lipidomic changes in head extracts from field-sampled queens (18% of annotated lipids were differentially abundant), but a comparatively small effect was observed in queens held in the laboratory (only 1% of annotated lipids were differentially abundant). We expected that a stronger effect would be observed in the more controlled laboratory environment; however, since a queen’s lipid intake is influenced by the workers’ royal jelly production, which is in turn influenced by the pollen the workers consume^50–52^, worker diet differences may have influenced the queen’s overall lipidome. This may also be linked to why such little variation in linolenic acid, which is provided to queens in royal jelly but is originally sourced from pollen^53^, was observed in the cage trial queens but not the field samples (**Figure 1**). Virus effects in the field may have also been exacerbated by interactions with other stressors or perhaps were only apparent when the queens exceeded a threshold level of ovary activation and metabolic demands (indeed, the ovaries of queens sampled from colonies were 2.3 times larger than those held in the laboratory; **Figure 2**). A consistent feature between the two datasets, though, is that triacylglycerol abundance declined with virus infection (**Figure 3**).

Triacylglycerols are one of the densest forms of stored biochemical energy, and this sweeping reduction indicates that infections are likely energetically demanding. This is consistent with the presence of our previously observed resource-allocation trade-off with immunity: For a trade-off to occur, a stimulus must first create a resource deficit, and this reduction in triacylglycerols may represent just that. However, extracts from the head are admittedly not the best tissue to analyze if one were to quantify energy reserves ― more relevant would be the fat body ― and future work should include comparative lipidomics analysis of multiple tissues to better relate pheromonal and physiological changes.

One advantage of conducting lipidomics on head extracts was that the head contains all known components of QRP^45^ and analyzing this tissue enabled comparisons with previously published data generated from small- and large-ovary queens^43^. We expected queens with small ovaries (laying restricted) to display a reduction in triacylglycerols relative to large-ovary (free laying) queens, mirroring the effect of virus infection (which is itself also an ovary-shrinking stimulus). However, this was not the case; few triacylglycerols (5.2% of all lipids) were identified in the small vs. large ovary dataset compared to the field samples (where triacylglycerols made up 12% of lipids). More striking was the fact that none of those that were differentially abundant were identified in both datasets, and among those in the small- vs. large-ovary dataset, some were upregulated as well as downregulated. Though direct comparisons may not be reliable due to generally poor dataset overlap, it appears that changes in triacylglycerols abundances in small- vs. large-ovary queens does not mirror the effect of virus infection. We speculate that this could be because laying restriction may not actually cause a resource deficit the way virus infection does. The restricted queens are likely fed less often by the workers, and their ovaries respond by shrinking, but they are also not laying eggs; therefore, resource inputs and outputs may still be balanced in this situation.

Most lipids declined with virus infection, but a select three increased: PGG2, a phospholipid (PEtOH 16:1/16:1), and a sphingolipid (SM 34:0;2O). The significance of the latter two is unclear, but PGG2 is interesting because it is an unstable lipid peroxide that is a precursor to other prostaglandins^54^, some of which release oviposition behavior in other insects^55^. We did not measure egg laying in the colonies these queens were heading, so we cannot confirm the expected inverse relationship between PGG2 and oviposition (and PGG2 was not identified in the dataset of small- and large-ovary queens), but it is an intriguing possibility. PGG2 was not upregulated in virus-infected queens in our cage trial (**Figure 3A**), so, if products of its conversion are important for oviposition, they appear only to be relevant in the colony context. PGG2 accumulation also suggests that oxidative stress may be, surprisingly, a collateral cost of laying reduction.

However, this idea is at odds with the fact that the majority of phospholipids downregulated in small-ovary queens contain polyunsaturated fatty acid (PUFA) residues (**Figure S3**), which are susceptible to oxidative attack. Therefore, based on PUFA membrane lipid abundances alone, small-ovary, non-laying queens should be under reduced oxidative stress than large-ovary, laying queens. Laying queens also likely have higher metabolic rates, which would increase reactive oxygen species generation. The overall low frequency of PUFA membrane lipids and low peroxidizability in queens relative to workers has been suggested as one factor enabling queens’ extraordinarily long lives compared to their non-reproductive daughters^56^, but the relative importance of PGG2 accumulation vs. reduced PUFA membrane lipid abundance is not yet clear.

Hemolymph proteomics analysis of the small- and large-ovary queens revealed that canonical immune proteins (antimicrobial peptides, lysozyme, and phenoloxidase activating factor) remain unchanged by ovary manipulation (**Figure 5A**). This indicates that the reproduction-immunity trade-off does not function in reverse (*i.e*., restricting ovary investment is not sufficient to elevate constitutive immune investment). However, ovary restriction did elevate two unique ferritin proteins (**Figure S4**), which are thought to reduce oxidative stress by sequestering iron cations (and thus preventing the formation of hydroxyl radicals^57^) as well as improve immunity by making iron unavailable to pathogens (reviewed in Pham and Winzerling^58^) ― roles that are supported by work in bumble bees^59^. Whether their main function here is in immunity, mitigating oxidative stress, or both is unknown, but it is noteworthy that these two unique ferritin proteins are upregulated in lockstep in small-ovary queens.

Regardless of the role of these ferritin proteins, ApoLP-III ostensibly does not switch from lipid trafficking to being an immune protein inducer when queens are in an ovary-restricted state (which would enable a bi-directional reproduction-immunity trade-off), as we originally hypothesized. Our previous rationale was that the reduced lipid demands of small ovaries could make more free ApoLP-III available to bind PAMPs and elevate immune effectors downstream of the Toll/Imd signalling pathways, assuming equivalent PAMP abundances. What we observed instead was that circulating immune effector abundance remained the same while ApoLP-III abundance decreased with ovary reduction (**Figure 5**). Therefore, it is still possible that our proposed mechanism is correct, but the immune induction capacity of ApoLP-III is attenuated by downregulating its expression to avoid investing in immune protein expression without an imminent fitness benefit. This downregulation of ApoLP-III may be precisely what is preventing a reverse reproduction-immunity trade-off from occurring. In a system in which ApoLP-III levels remain unchanged and PAMPs are introduced, as was observed upon virus infection previously^25^, ApoLP-III could still be serving as a switch that governs reproduction and immunity investment. We have yet to reconcile the observation that ApoLP-III was not induced by experimental virus infection with the results of prior queen survey data showing that ApoLP-III was more abundant in failing (often virus-infected) queens compared to healthy queens, while ApoLP-I/II was less abundant^24^. These lipophorins are clearly tied to queen health and ovary investment but the mechanisms cannot be fully defined without gene knockdown studies or factorially designed experiments with infected and uninfected, caged and uncaged queens. Our ovary section and oocyte protein data do support that ApoLP-III, ApoLP-I/I, and vitellogenin transport lipids in a manner consistent with what is described in other insects^11,15,21^, though, which forms the foundation for further experimentation.

Importantly, in our previous study on the relationship between queen pheromone profiles and ovary size^43^, we reported that the abundance of methyl oleate remained unchanged based on metabolomics-derived peak areas. However, lipidomics data obtained from the same samples through a two-phase extraction demonstrate a clear correlation between methyl oleate abundance and ovary size. This discrepancy likely arises from the presence of co-eluting isomers in the metabolomics dataset, which affected the accurate determination of methyl oleate, despite the use of external standards. While the quantification of other compounds in the original metabolomics dataset is not affected, the lipidomics approach used here offers isomeric resolution for analyzing methyl oleate and several hydroxylated fatty acids. This aligns with its established role in queen reproductive signaling and highlights the importance of analytical specificity when analyzing complex lipid mixtures. This serves as a cautionary tale for lipid analysis: many lipid isomers, particularly those with subtle structural differences (*e.g.,* double-bond positioning or branching), are not routinely resolved by shotgun lipidomics or traditional methods^60^. These isomers are often absent from literature databases and may co-elute in standard workflows, leading to misinterpretations of lipid abundance and function. Our outcomes highlight the necessity of isomer-sensitive techniques when studying lipids in complex biological systems. Overlooking these nuances risks missing critical biological insights, especially in contexts where lipid structural diversity drives physiological outcomes.

## Conclusion

In this body of work our goal was to answer five core questions: 1) does virus infection reduce abundance of QRP components, 2) what other lipids are affected by virus infection, 3) is ovary restriction alone sufficient to induce the same changes to pheromones and lipids, 4) does ovary restriction cause a reverse reproduction-immunity trade-off (*i.e.*, are immune effectors elevated under ovary restriction), and 5) could ApoLP-III be mediating the reproduction-immunity trade-off induced by virus infection? We have answered all these questions with varying degrees of certainty. Virus infection reduces methyl oleate abundance as well as many triacylglycerols, indicating a depletion of stored energy. Since ovary restriction also reduces methyl oleate abundance (but not triacylglycerols), the impact of virus infection on methyl oleate appears to be an indirect effect of diminished ovaries. Since canonical immune proteins do not increase under ovary restriction, the reproduction-immunity trade off appears to only operate under immune stimulation and not restricted laying. Our data are consistent with positioning ApoLP-III as a candidate molecular switch that could govern reproduction vs. immunity investment, but more detailed experimentation is necessary to assert this claim. Future work should also focus on determining if a decline in methyl oleate is sufficient to trigger supersedure cell rearing in colonies, as the mechanism of initiating this behavior is unknown.

## Methods

### Cage trial and field sample queens

The samples described here were derived entirely from cage trials and field samples that have been previously described^25^. Briefly, in the cage trial, young (< 1 month old) queens were microinjected with either saline, a live virus inoculum (a mix of 100 million BQCV copies and 2 million DWV-B copies, to approximate common co-occurrence levels of these viruses in queens), or a UV-inactivated virus inoculum (the same concentration of virus particles as the live inoculum that were inactivated by UV-crosslinking), with n = 9 queens in each group. The queens were housed in queen monitoring cages^61^ with ∼100 young workers as well as *ad libitum* fondant, pollen substitute, and water. The cage walls were made of replaceable egg-laying plates to facilitate laying. The queens were observed for seven days after injection to monitor egg laying, then were euthanized, dissected, and samples were stored at −70 °C. See Chapman *et al.*^25^ for complete experimental details, including RT-qPCR viral analyses.

The field samples are derived from age-matched queens (from the same source as the cage trial) heading 32 five-frame nucleus colonies, as previously described^25^. Colonies were located in two different apiaries (sites) in a balanced design, and all colonies received *ad libitum* pollen substitute (15% pollen patties, Global) and Hive Alive fondant for the duration of the experiment. The queens were blindly divided into two groups (n = 16 each), with one receiving a saline injection and the other receiving the same live virus inoculum described above. However, upon completion of the trial (seven weeks after injection) RT-qPCR analysis showed that the queens receiving the live virus inoculum did not have higher viral loads than the saline controls; rather, the queens’ viral abundances correlated significantly with worker viral abundance (**Figure S1**). We, therefore, decided to treat the experiment as an observational dataset, correlating queen virus abundance to traits of interest irrespective of the experimental group. Queens were euthanized and stored as above. See Chapman *et al.*^25^ for complete experimental details, including viral analyses.

### Queens sampled for ovary section analysis

To compare lipid transporter abundance in ovary sections, we dissected queens acquired from a routine apiary requeening sweep and divided ovaries into three parts with a scalpel as illustrated in **Figure S5**. Queens originated from a single batch of Northern Californian imports that headed colonies and overwintered in a single apiary, then were allowed to grow in size naturally through the late winter and early spring. Queens were then sampled in April, transported to the laboratory in queen cages with 5 attendant workers and queen candy, then anesthetized (5 min carbon dioxide exposure), decapitated, and dissected within two hours of being removed from colonies. Ovaries were divided into sections with under a dissection microscope and mature oocytes were delicately removed from the lateral oviducts, then proteins were extracted from tissues exactly as previously described^62^.

### Lipid extraction

Lipids (including QRP components) were extracted from cage trial and field sample queen heads using methods exactly as previously described^43^. Heads were chosen because this is the only body segment known to contain all QRP compounds^45^, and this tissue enables comparisons to previously published data^43^. Briefly, we followed a two-phase extraction procedure^63^. The queen heads were homogenized in methanol extraction solvent (400 µl 75% methanol, 0.1% butylated hydroxytoluene, in ddH_2_O), then methylated tert-butyl ether (1 mL) was added and the samples were incubated for 1 h (1000 rpm shaking, room temperature). Next, clarified extract was mixed with water to induce phase separation. After a brief incubation at room temperature (10 min), samples were centrifuged (14,000 g, 15 min, 4°C) and 250 µl of the upper phase (lipids) was retained for lipidomics. We used a combination of internal and external standards to facilitate peak identification (see McAfee *et al*.^43^ for complete details). The samples were stored at −70 °C in solution until LC-MSMS analysis.

### Lipidomics LC-MS/MS analysis

Lipidomics analysis was conducted as previously described [32]. Lipid extracts were resuspended in a solution of 70% acetonitrile and 30% isopropanol, and separated using a Vanquish UHPLC system (Thermo) equipped with an ACQUITY UPLC CSH C18 analytical column (130 Å, 1.7 µm, 2.1 mm × 100 mm, Waters). A multi-step gradient was employed, increasing from 20% to 99% mobile phase B over 18 minutes. Mobile phase A consisted of water, while mobile phase B was a mixture of acetonitrile and isopropanol, both containing 10 mM ammonium formate and 0.1% formic acid. The column was maintained at a temperature of 65°C with a flow rate of 0.4 mL/min, and the samples were kept at 4°C. Mass spectrometry (MS) analysis was performed on a Bruker Impact II QTOF in both positive (ESI+) and negative (ESI−) electrospray ionization modes. The capillary voltages were set to +4,500 V for ESI+ and - 3,800 V for ESI−, with a dry gas temperature of 220°C, and a scan range of 100–1,700 m/z. To enhance sensitivity for hydroxy fatty acids and improve structural coverage, additional improvements were implemented for the ESI+ dataset: the spectral acquisition rate was set to 2 Hz, and a stepped MS/MS collision energy varying from 100% to 250% was applied, with 50% timing at each energy level. Dynamic MS/MS acquisition was utilized, targeting an intensity of 1125 counts, with an acquisition frequency ranging from a minimum of 20 Hz to a maximum of 40 Hz. Finally, structural information was improved by implementing a 15-second active exclusion window, minimizing redundant fragmentation and improving isomer coverage.

All mass spectrometry data are housed at Metabolomics Workbench (www.metabolomicsworkbench.org). Data derived from cage trial and field sample queens were newly acquired (doi:10.21228/M88838; project PR002316), whereas data from ovary-restricted queens were previously published (doi:10.21228/M81B11; project PR001924)^43^.

### Lipidomics data processing

Annotations were conducted following the Metabolomics Standards Initiative (MSI) reporting criteria, as previously detailed^64^. The raw data were processed using Progenesis QI software (version 3.0.7600.27622) with the METLIN plugin (version 1.0.7642.33805, NonLinear Dynamics). This process included peak picking, alignment, deconvolution, normalization, and database querying. Level 2 annotations were assigned by screening features against databases such as METLIN^65^, GNPS^66^, HMDB^67^, MassBank of North America^68^, and LipidBlast^69^. For compounds detected in both ion modes, the compound with lower variability in quality controls (QCs) was retained. Detailed information on the queens from which the previously published ovary restriction dataset was derived has been described elsewhere^43^. Briefly, queens were reared simultaneously at a single queen production operation from a single mother colony. Queen cells were placed in mating nucleus colonies and once emerged, the queens were allowed to freely mate. Two weeks after emergence, a cohort of mated queens were placed in JZBZ queen cages, which are large enough to house the queen but do not support egg laying, and placed in a “queen bank” (a large colony with sufficient young bees to feed the queens through the cage screen). One month after emergence, n = 10 of these queens were sampled, making up the “small ovary” (caged) group. Another n = 10 queens that had remained laying in their nucleus colonies were sampled on the same date, making up the “large ovary” (uncaged) group. Annotated lipid abundance data were obtained from head extracts exactly as previously described^43^ and are also available in **Supplementary Data 1**.

### Statistical analysis of lipidomics data

Peak areas corresponding to the quantifiable components of the queen retinue pheromone (9-ODA, 9(R)-HDA, HVA, HOB, LEA, MO, and CA) were identified with high confidence by comparing spectra and retention times to external standards. These *a priori* compounds of interest were analyzed separately from the complete lipidomics dataset. We lacked an external standard for 9(S)-HDA; therefore, only the R enantiomer was evaluated. PA (palmityl alcohol), which is also a QRP component^45^, occurred below our limit of detection. This means that seven of the nine QRP components were assessed here. See McAfee *et al*.^43^ for vendor sources of external standards.

We used R (version 4.3.0) via R Studio (version 2023.09.1+494) for all subsequent statistical analyses^70^. Queen and colony metadata (virus abundances, ovary masses, presence of supersedure cells, *etc.*) are publicly available from our previous publication^25^. We modelled each of the seven QRP components individually using either additive queen virus load or ovary mass as a predictor in a simple linear model. Appropriateness of model fit was confirmed by inspecting residual distributions. A Bonferroni correction (α = 0.05/7 = 0.0071) was applied.

In our previously published results^25^, we reported that queen total viral load was a negative but not significant predictor of ovary mass, and that total viral load was a marginally non-significant predictor of supersedure cells; however, there were inconsistencies in the way that total viral load was calculated, and in both cases the relationships are actually significant. In the prior study, copy numbers (per ng of RNA) were first transformed (natural logarithm) then summed, but this approach biases the total viral load metric by weighting queens infected with multiple viruses more heavily, instead of reflecting the simply summed copy numbers, as we have previously done^24,71^. To correct this error and regain consistency with our previous studies, we recalculated total viral load by first summing copy numbers per ng RNA, then log10 transforming, and re-analyzed worker viral load as a predictor of queen viral load (linear model), queen viral load as a predictor of ovary mass (linear model), and queen viral load as a predictor of supersedure cell presence (generalized linear model). The results of these reanalyses are shown in **Figure S1**. In the present study, we use the corrected method of calculating total viral load in all instances.

To assess lipidomic changes, we scaled and combined all annotated lipids identified in positive and negative ion mode into a single dataset of 336 compounds. We log10 transformed the data and visually confirmed normality of the overall data distribution. We used tools within the limma package^72^, which enables empirical Bayes variance estimation, to identify significant relationships between lipid compound abundances and total virus load (continuous predictor) for both the cage trial (final n = 27) and field sample data (n = 29, down from 32 due to three instances of queen rejection, supersedure, or mishandling). The field sample lipidomics data were additionally analyzed with respect to the presence or absence of supersedure cells in the colony (categorical predictor, two levels), and, separately, ovary mass (continuous predictor). In all cases, false discovery rates were controlled to 5% using the Benjamini-Hochberg method. Because total virus load predicts both ovary mass and the presence/absence of supersedure cells, including both variables in the same model would obscure our ability to detect potentially meaningful relationships. Therefore, we instead analyzed the data with respect to each variable separately and compared the overlap among significantly different lipids. To compare abundance of specific lipids between two groups, we used simple linear models and inspected residual distributions to confirm appropriateness of fit.

### Structural enrichment of lipid classes

We used ChemRich^73^ to detect statistical enrichment of lipid structural classes with respect to p values associated with total viral load. We obtained CIDs, SMILES, and InChiKeys for each lipid compound from the PubChem database (https://pubchem.ncbi.nlm.nih.gov/). For instances where the positions of double bonds or functional groups were unknown, we assumed the compound was one of the available isomers not already represented in the dataset. This approach was justified because enrichment analysis of structural classes depends on the presence, rather than the specific positions, of functional groups or unsaturation; therefore, these ambiguous isomers are best handled by still representing them in the data. Significance of enrichment was controlled to 5% FDR using the Benjamini-Hochberg method.

### Proteomics data acquisition and data processing

To assess the impact of ovary restriction on immune effector and lipid transporter expression, we analyzed publicly available hemolymph proteomics data from ovary restricted and unrestricted queens^48^ housed on the MassIVE proteomics data archive (https://massive.ucsd.edu/ProteoSAFe/static/massive.jsp) under accession MSV000094277. These data were partially analyzed in the associated publication, but data from the caged queen cohort were not presented as we were previously focused on assessing queen aging in their natural (unrestricted) environment. All samples were processed in parallel with those previously described and the data acquisition and processing methods are identical^48^. These are the same ovary restricted queens as described in the lipid analysis section (lipids were extracted from their heads and proteins were extracted from their hemolymph.

Proteomics data for the ovary section samples were acquired by injecting 250 ng on an Easy nLC-1000 system (Thermo) connected in-line to an Impact II QTOF mass spectrometer (Bruker). Liquid chromatography and data acquisition methods were exactly as previously described^74^. Raw data were searched using MaxQuant^75^ version 1.6.1.0 with default search options except that match between runs was enabled. The data were searched against the honey bee reference proteome available on NCBI (based on the build HAv3.1, downloaded Nov 18th, 2019) with all honey bee virus and *Nosema* (*Vairimorpha*) proteins included in the FASTA file. The protein quantification data are supplied in **Supplementary Data 1.**

### Statistical analysis of proteomic data

Proteome-wide differential expression analysis of the small- and large-ovary queen hemolymph was conducted using the limma package^72^ essentially as described for the lipidomics data analysis. The exceptions are that the data were first filtered to remove contaminant protein sequences and proteins identified in fewer than 50% of the samples. FDR was controlled to 5% using the Benjamini-Hochberg correction. Specific proteins of interest (vitellogenin, ApoLP-III, and ApoLP-I/II) were further analyzed using a simple linear model to compare abundance between two groups (small-ovary vs. large-ovary queens). The abundance of these proteins in the independent ovary section dataset were assessed using linear mixed models to compare abundance in ovary sections (four levels: germarium, early vitellariums, late vitellarium, and germarium). The mixed model also included queen as a random effect (10 levels) to account for non-independence of samples.

## Acknowledgements

We would like to acknowledge the mass spectrometry core facility team ― Jason Rogalski, Renata Moravcova, and Jeanne Yuan ― for their technical expertise and role in the proteomic data acquisition.

## Author contributions

AM wrote the first draft of the manuscript, conducted statistical analysis, and generated the figures. AC designed and conducted the cage experiment. AM designed and conducted the field analysis with assistance from AC. AM prepared the pheromone and lipidomics samples, and AAM acquired the mass spectrometry data. SEH generated the caged and uncaged queens. LJF and SEH acquired funding and provided resources to support this research. KM and DRT advised AM and made intellectual contributions to the analysis and writing. All authors edited the manuscript and approved of the submitted version.

## Supplemental information

Document S1. Figures S1-S5.

Data S1. Lipidomics data, metadata, statistical results, and chemrich summaries.

